# Probabilistic forward replay of anticipated stimulus sequences in human primary visual cortex and hippocampus

**DOI:** 10.1101/2022.01.26.477907

**Authors:** Matthias Ekman, Giulia Gennari, Floris P. de Lange

## Abstract

The ability to recognize and predict future spatiotemporal sequences is vital for perception. It has been proposed that the brain makes ‘intelligent guesses’ about future inputs by forward replaying these events. However, it is unknown whether and how this mechanism incorporates the probabilistic structure that is inherent to naturalistic environments. Here we tested forward replay in human V1 and hippocampus using a probabilistic cueing paradigm. Participants were exposed to two visual moving dot sequences (A and B) that shared the same starting point. Each stimulus sequence was paired with either a high or a low tone that predicted which sequence would follow with 80% cue validity (probabilistic context) or 50% cue validity (random context). We found that after exposure, the auditory cue together with the starting point triggered simultaneous forward replay of both the likely (A) and the less likely (B) stimulus sequence. Crucially, forward replay preserved the probabilistic relationship of the environment, such that the likely sequence was associated with greater anticipatory V1 activity compared to the less likely stimulus sequence. Analogous to V1, forward replay in hippocampus was also found to preserve the probabilistic cue-sequence relationship. Further, the anterior hippocampus was found to represent the predicted stimulus sequence, irrespective of the input, while the posterior hippocampus revealed a prediction error-like signal that was only observed when predictions were violated. These findings show how mnemonic and sensory areas coordinate predictive representations in probabilistic contexts to improve perceptual processing.

## Introduction

There is mounting evidence that visual computations are inherently predictive (Rust & Palmer, 2021; Summerfield & De Lange, 2014) and rely on past experiences to anticipate future events (Gavornik & Bear, 2014; Xu et al., 2012). We previously found that merely presenting the starting point of a moving dot sequence triggered an activity wave in human primary visual cortex (V1) that resembled the full stimulus sequence (Ekman et al., 2017). This anticipatory activity wave, encoding possible future trajectories, is sometimes referred to as forward replay (Carr et al., 2011; Derdikman & Moser, 2010; Diba & Buzsáki, 2007), and is thought to play a crucial role in learning (Jadhav et al., 2012) and planning (Buzsáki & Moser, 2013; Momennejad, 2020; Pfeiffer & Foster, 2013; Wikenheiser & Redish, 2015).

While simple predictions based on extrapolation of motion trajectories could be implemented locally within V1 (Alink et al., 2010; Lee & Mumford, 2003; Rao & Ballard, 1999), it has been suggested that more complex relationships like context-modulation (McClelland et al., 1995), or cross-modal associations (Kok & Turk-Browne, 2018), depend on the memory system, specifically the hippocampus, for driving memory-based expectations in sensory areas like V1 (Hindy et al., 2016; Ji & Wilson, 2007). Supporting this notion, a recent study has shown that predictive sequence representations in V1 became absent after hippocampus lesioning (Finnie et al., 2021).

Previous studies have shown that hippocampus acquires regularities of both simple (Aitken & Kok, 2021; Finnie et al., 2021; Kok & Turk-Browne, 2018; Schapiro et al., 2012) and more complex stimulus sequences (Kurth-Nelson et al., 2016; Liu et al., 2019; Schapiro et al., 2013; Schuck & Niv, 2019), that can be exploited to encode representations of future trajectories (Diba & Buzsáki, 2007; Dragoi & Tonegawa, 2011) and potentially be shared via feedback connections with sensory cortices like V1 (Finnie et al., 2021; Hindy et al., 2016; Kok & Turk-Browne, 2018).

However, natural environments are often inherently probabilistic and present us with the uncertainty of multiple, often competing, future sequences. It remains an open question how probabilistic future trajectories are represented after learning. One possibility is that hippocampus and V1 encode representations of multiple future states simultaneously. Alternatively, the hippocampus could simultaneously represent multiple options, but only communicate the most likely (or relevant) future outcome to downstream sensory area V1.

To answer this question, we conducted an fMRI study where participants were presented with a probabilistic cueing paradigm, in which an auditory cue (tone A or B) was followed by one of two moving dot sequences (sequence A or B). After learning, we introduced occasional omission trials, where only the cue and the starting point of the sequence were presented, while the rest of the visual sequence was omitted. These omission trials allowed us to study expectations of future stimulus sequences in the absence of physical stimulation. Further, by systematically varying the cue-sequence probability across two separate sessions, we were able to test whether anticipatory activity in hippocampus and V1 also encodes the probability of the anticipated sequences.

To preview our results, we found that only presenting the cue and sequence starting point triggered V1 BOLD activity that reactivated both sequences A and B, with stronger reactivation of the sequence that was more likely given the cue. Importantly, we observed a spatiotemporal dissociation in hippocampal representations. Predictive representations emerged temporally early in the anterior hippocampus and were coordinated with V1, while posterior hippocampus showed a later, prediction error-like response, in case predictions were violated. The finding that V1 and anterior hippocampus activity patterns preferentially represent the likely sequence suggests that sensory and mnemonic representations take the probabilistic relationship of the environment into account when predicting future trajectories.

## Results

Human observers (N=28) were exposed to a probabilistic cueing paradigm, in which an auditory cue (cue A: high tone, or cue B: low tone) was followed by one of two possible visual dot sequences (Sequence A or Sequence B; **Figure 1A**). Note that both sequences share the identical starting point location. Therefore, merely presenting the starting point does not provide any information about the respective sequence.

**Figure 1.**
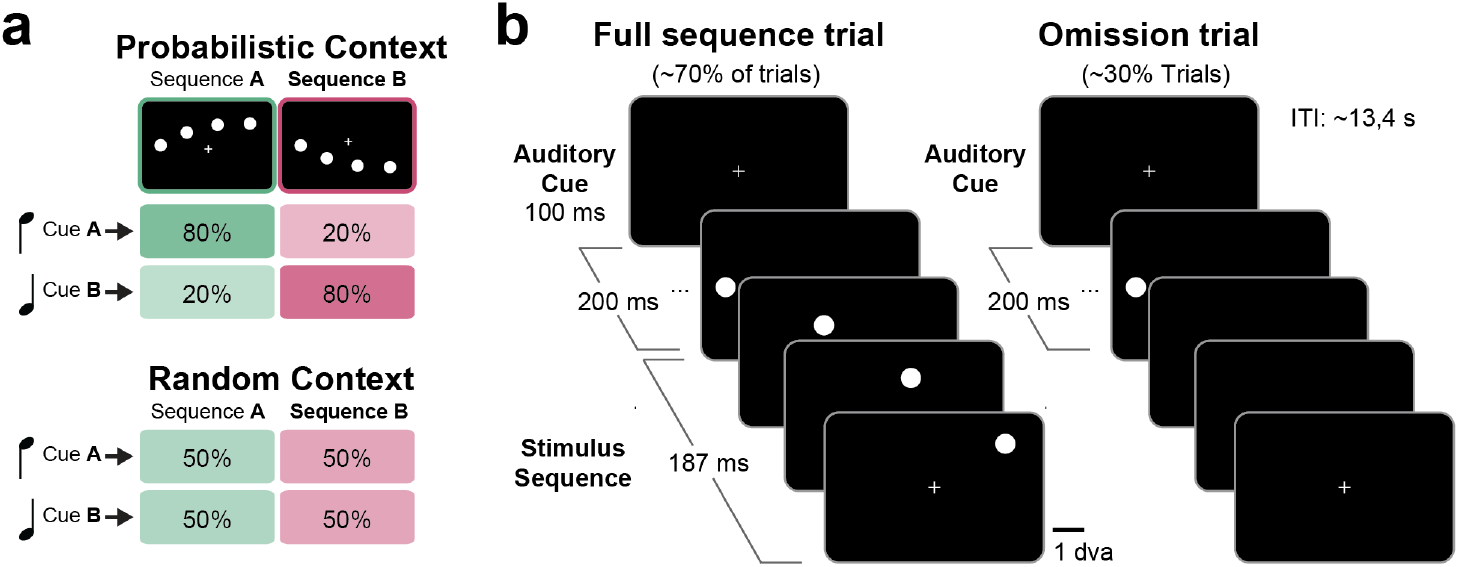
Probabilistic cueing paradigm. (**a**) Probabilistic cue contingency and context modulation. In the probabilistic context, Cue A was followed by sequence A 80% of the time and sequence B 20% of the time (top). In the random context, the cue contingency was at chance level (50%). (**b**) Stimulus timing for full sequence and starting point only trials. An auditory cue (high, low) was followed by one of two possible moving dot sequences consisting of four dots. During omission trials, only the auditory cue together with the first dot of the sequence was presented. During an initial learning period, only full sequence trials were shown and participants were instructed to detect a small temporal delay in the presentation of the last dot of the sequence. After learning, participants were randomly presented with full sequence trials and starting point trials, while performing a detection task at fixation. Dva, degrees of visual angle.

After an initial exposure period (~12 min) with full sequence trials only, we continued to present full stimulus sequence trials (full sequence condition), but occasionally only the auditory cue together with the starting point of the sequence was shown, omitting the remaining sequence dots (omission condition) (**Figure 1B**).

We hypothesized that V1 may forward replay the remaining sequence dots and that cortical activity during cue-triggered replay should resemble the activity of the full sequence condition at retinotopically defined sequence locations. In order to probe the potential probabilistic representation of forward replay, we manipulated the probabilistic cue-sequence association in the following way. In the probabilistic context, the auditory cue A was followed by sequence A 80% and sequence B 20% of the time (**Figure 1A**). Conversely, cue B was followed by sequence B 80% of the time. In the random context, the cue validity was at 50% (chance level) for cue A and cue B. In other words, in the random context there was no probabilistic relationship between the auditory cue and the visual sequence, as both sequences were equally likely to be followed by either cue A or B. For the context manipulation, participants were tested in two separate and counterbalanced sessions (probabilistic vs. random context) that were on average 14 days apart.

### Internalization of the probabilistic cue-sequence structure leads to behavioral benefits

First, we tested whether participants indeed learned the probabilistic cue-sequence relationship. During the initial exposure, participants were instructed to maintain fixation while detecting an infrequent temporal delay in the visual sequence (in 15% of trials the last dot of the sequence was delayed by 170 ms). We reasoned that being able to correctly predict the upcoming sequence A vs B should facilitate the detection of the temporal delay. We therefore predicted that if participants successfully learned the cue-sequence relationship, behavioral responses in the probabilistic context should be faster for cue-valid trials (e.g. sequence A was cued and presented), compared to cue-invalid trials. Conversely, no benefit should be present in the random context. Note that due to the chance-level cue contingency the random context does technically not have valid, or invalid trials. In this case we labeled trials as valid or invalid based on the cue-association in the probabilistic context.

As expected, reaction times (RTs) in valid trials were found to be significantly faster (t(27) = −3.03, P = 0.005) compared to RTs in invalid trials (RT_valid_ = 459 ms; RT_invalid_ = 471 ms), confirming that subjects built an internal representation of the probabilistic cue-sequence relationship. Further, in line with our prediction, no behavioral facilitation was found in the random context (t(26) = 1.49, P = 0.15; RT_valid_ = 462 ms; RT_invalid_ = 458 ms). The Cue [Valid, Invalid] x Context [Probabilistic, Random] interaction was found to be significant (F(1,26) = 9.86, p = 0.004, η^2^ = 0.03), indicating a significant context modulation of participants’ performance.

### Probabilistic forward replay in early visual cortex V1

Probing BOLD activation patterns in early visual cortex (V1), we first wanted to confirm that the physical presentation of the visual sequence elicits a sequence-specific activation pattern at the receptive field locations that receive bottom-up visual input. To this end, we first determined V1 receptive fields (RFs) in a separate session using population receptive field (pRF) mapping (*see Materials and Methods*). Then we compared BOLD activity at V1 RFs covering the last three sequence locations during presentation of sequence A and sequence B. As expected, presentation of sequence A elicited higher BOLD activity at RFs corresponding to sequence A compared to RFs corresponding to sequence B (**Figure 2A**; paired-sample t-test for BOLD averaged across locations, sequence A: t(27) = 10.80, p = 2.65 × 10^−11^). The reverse BOLD pattern was observed when sequence B was presented, i.e., higher BOLD activity at sequence B RFs compared to sequence A RFs (t-test t(27) = 7.39, p = 5.87 × 10^−8^). BOLD activity at the non-stimulated sequence RFs (i.e., sequence B RFs during cue A/sequence A trials) was not significantly different from baseline activity (one-sample t-test for BOLD averaged across locations and sequence A/B: t(27) = −0.98, p = 0.34).

**Figure 2.**
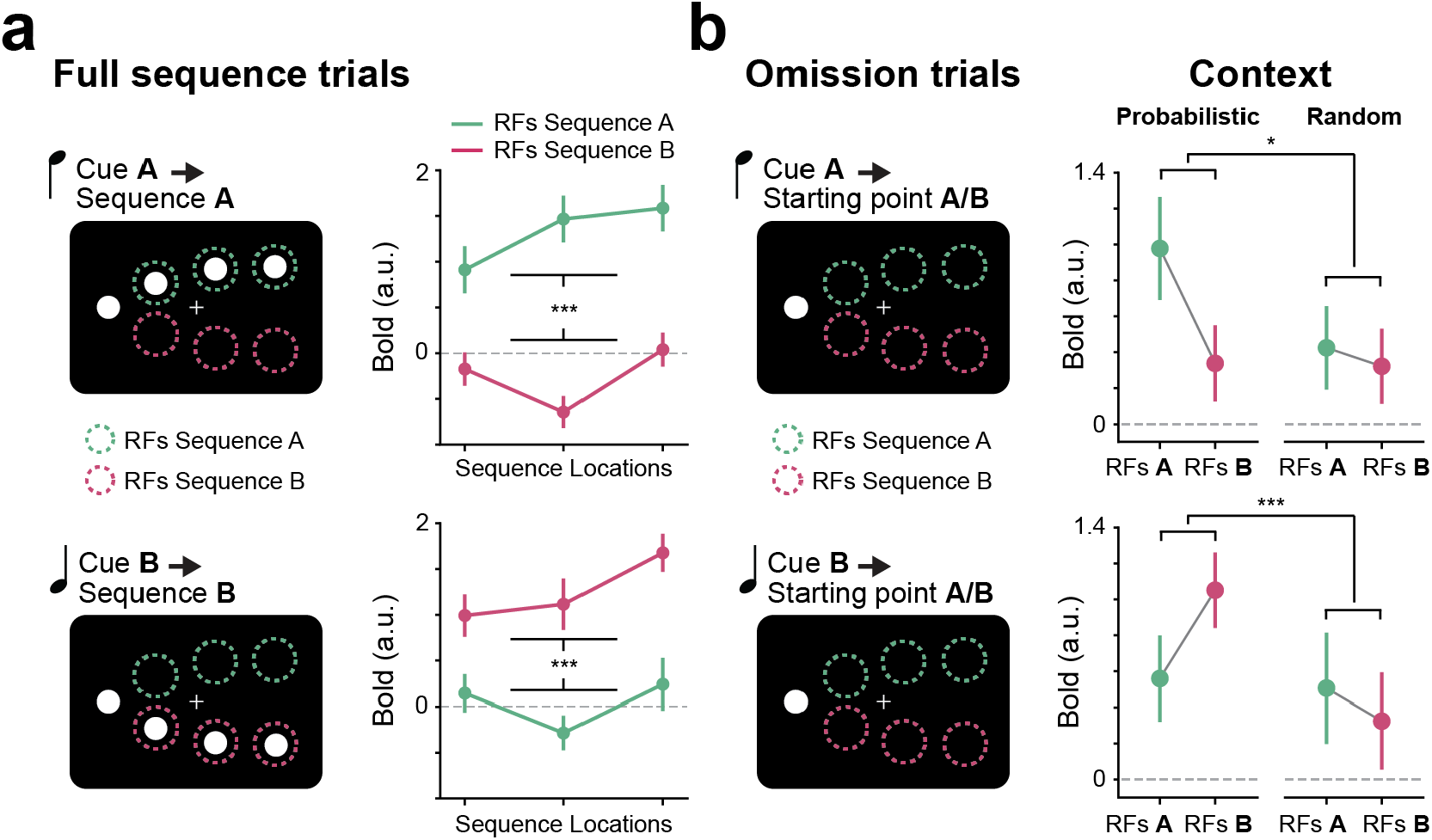
Probabilistic forward replay in human V1. (**a**). BOLD activity during full sequence trials for cue A → sequence A trials (top) shows greater BOLD activity at sequence A receptive field locations (RFs, green) compared to sequence B RFs (red). The reverse pattern was observed for cue B → sequence B trials (bottom), i.e. greater BOLD activity at sequence B RFs (red) compared to sequence A RFs (green). (**b**). Anticipatory BOLD activity during omission trials for cue A → starting point A/B (top) shows greater BOLD activity at averaged sequence A RFs compared to sequence B RFs in the probabilistic context, but not in the random context. The reverse pattern was observed for cue B replay trials (bottom). Error bars denote ± s.e.m.; ***P<0.001; *P<0.05.

After learning, participants were presented with full sequence trials, but occasionally only the auditory cue together with the starting point of the sequence was shown (**Figure 2B**). We examined whether the sequence-specific activation pattern that was observed during full sequence trials was re-instantiated in response to omission trials where only the sequence starting-point was presented. Importantly, we predicted that the forward replay of the sequence would be modulated by the probabilistic cue-sequence relationship, such that cue A would elicit higher BOLD activity at the RFs overlying sequence A, while cue B would elicit higher BOLD activity at the RFs overlying sequence B. Additionally, if forward replay took the probabilistic cue-sequence relationship into account, the preferable replay of sequence A vs B should only be present in the probabilistic context, while replay in the random learning context should represent both sequences to an equal amount.

In line with our predictions, V1 BOLD activity during omission trials indeed revealed a preferential re-activation of the likely sequence RFs compared to the less likely sequence RFs (i.e. greater activity of sequence A vs B RFs in cue A trials) in the probabilistic context (paired sample t-test t(27) = 3.88, p = 0.001), but not in the random context (t(26) = 0.47, p = 0.64), culminating in a significant Location [RFs A, RFs B] x Context [Probabilistic, Random] interaction (F(1,26) = 10.32, p = 0.003, η^2^ = 0.04). BOLD activity in the probabilistic context was significantly higher than baseline for the likely sequence RFs (paired sample t-test, t(27) = 6.51, p = 5.532 × 10^−7^) and for the less likely sequence RFs (paired sample t-test, t(27) = 2.70, p = 0.012). Similarly, in the random context both (equally likely) sequence RFs showed significantly greater BOLD activity compared to baseline for cue A trials (paired sample t-test, t(26) = 2.49, p = 0.019) and cue B trials (paired sample t-test, t(26) = 2.28, p = 0.031).

### Coordinated forward replay in V1 and hippocampus

Next, we investigated whether the probabilistic forward replay that we observed in V1 was also present in the hippocampus. Note that while the hippocampus formation and nearby entorhinal cortex might feature a coarse representation of visual space (Killian et al., 2012; Knapen, 2021; Nau et al., 2018; Silson et al., 2020), it does not feature the same fine-scaled retinotopic organization present in V1 (Dumoulin & Wandell, 2008). Therefore, we employed a different analysis approach for hippocampus, focusing on population activity pattern across the entire ROI. In order to probe hippocampal representations, we trained a multivariate pattern classifier to distinguish between sequence A and B representations during full sequence trials and then applied the classifier to omission trials (*see Materials and Methods*). For comparison sake we report the classifier results for both hippocampus and V1. As expected, the classifier results for V1 mirror what was already reported based on the BOLD activity above, namely a preferential representation of the likely sequence in the probabilistic context, i.e., representation sequence A > B during cue A trials and representation sequence A < B during cue B trials (**Figure 3A**; t(27) = 2.270, p = 0.031) and no preferential representation in the random context (t(26) = 0.135, p = 0.894).

**Figure 3.**
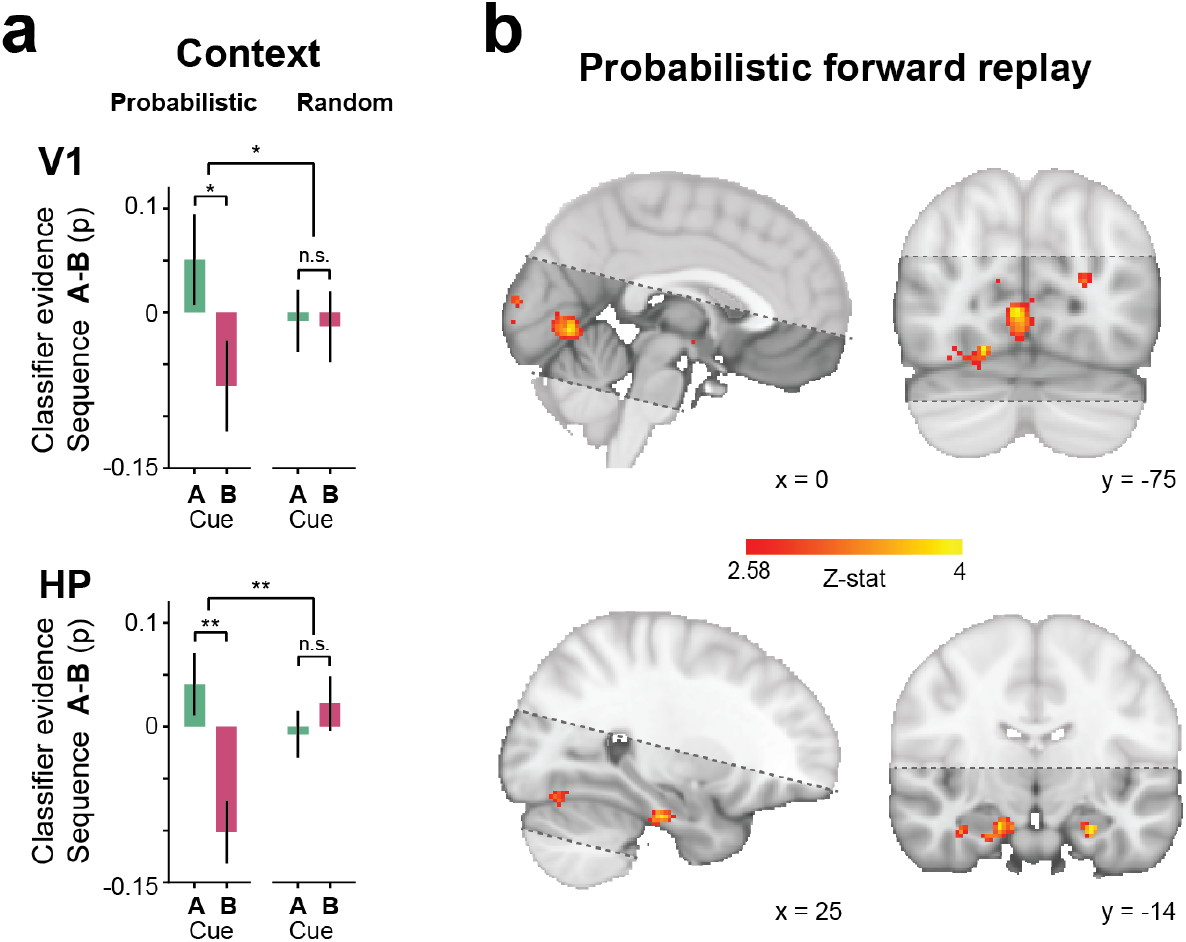
Probabilistic forward replay in V1 and hippocampus. (**a**). A classifier was trained to distinguish between full sequence A and sequence B presentations and then applied to replay trials. In the probabilistic context, classifier evidence (e.g., probability of sequence A) showed a biased representation toward the more likely sequence (e.g., evidence sequence A > B when cue A was presented) in both V1 (top) and hippocampus (HP, bottom). In the random context, when both sequences were equally likely, classifier evidence showed no biased representation toward either sequence. (**b**). Group averaged searchlight decoding analysis revealed that the decoding results are localized primarily in the early visual cortex and the anterior hippocampus (threshold at z=2.58 uncorrected, to visualize the extent of the spatial localization). The black dashed lines denote the approximate MRI coverage. Error bars denote ± s.e.m.; \*\**P*<0.01; \**P*<0.05.

Importantly, results for the hippocampus revealed a similar pattern to what was observed in V1. Specifically, sequence representation in hippocampus was greater for the likely sequence compared to the less likely sequence in the probabilistic context, i.e., representation sequence A > B during cue A trials and representation sequence A < B during cue B trials (**Figure 3A**; t(27) = 3.464, p = 0.002). In contrast, no biased sequence representation was found in hippocampus when the two sequences were equally likely (t(26) = −0.749, p = 0.460). The Cue [A, B] x Context [Probabilistic, Random] interaction was found to be significant in V1 (F(1,26) = 5.43, p = 0.01, η^2^ = 0.003) and hippocampus (F(1,26) = 10.242, p = 0.004, η^2^ = 0.093).

We also examined whether the reported forward replay was present outside the predetermined V1 and hippocampus regions using a whole-brain searchlight analysis. Results of the whole-brain analysis indicate that, within our restricted field of view, the observed forward replay effects were selectively located in the anterior, bilateral hippocampus and early visual cortex (**Figure 3B**).

Given the simultaneous forward replay in V1 and hippocampus, the question arises whether the two regions coordinate their respective representations. To answer this question, we performed an across subject correlation analysis testing whether the biased sequence representation toward the likely sequence in hippocampus was related to the biased sequence representation in V1. We reasoned that correlated representations between V1 and hippocampus might indicate that representations were shared between the areas (Clarke et al., 2021). Our results revealed a significant V1-hippocampus correlation in the probabilistic context (Spearman r = .49, p = 0.008), but not in the random context (r = −.23, p = 0.25). The difference in correlation between the probabilistic and random context was found to be significant (p = 0.004).

### Anterior Hippocampus represents prediction also during invalid trials

Thus far we have shown that both V1 and hippocampus preferentially represent the presented (and expected) sequence over the not presented (and less expected) sequence during cue-valid full sequence trials.

Next, we investigated how representations changed when predictions were violated during cue-invalid trials (20% of the trials, e.g. cue A → sequence B; **Figure 3A**). One could expect that representations keep tracking the predicted sequence, or alternatively, representations could switch from the predicted to the physically presented sequence. While it has been well documented that V1 is modulated by prior expectations (Ekman et al., 2017; Kok et al., 2012; Summerfield & De Lange, 2014), V1 is more strongly driven by bottom-up stimulation. Therefore, we hypothesized that overall, V1 representations should be dominated by the physical stimulus and should therefore preferentially represent the shown (but less expected) sequence during cue-invalid trials. For hippocampus our hypothesis was less clear. A recent study has shown that hippocampus preferentially represents the predicted content, irrespective of the bottom-up visual input (Kok & Turk-Browne, 2018). However, hippocampus has also been involved in novelty detection (Kafkas & Montaldi, 2018) and representation of surprise (Bein et al., 2020) in case of violated predictions. In our decoding analysis such a prediction error-like response during invalid trials would manifest as a biased representation toward the presented sequence. Because our previous results for valid trials located the predictive relationship mainly in the anterior part of the hippocampus and given that recent studies point to a differential representations along the long axis of the hippocampus (Aitken & Kok, 2021; Collin et al., 2015; Silson et al., 2020), we carried out an additional exploratory analysis, dividing the hippocampus region into two separate subparts: an anterior and a posterior hippocampus region. Further, to dissociate predictive and stimulus related representations that should occur earlier in time from surprise and prediction error-like representations that one would expect to occur later in time, we used a time-resolved decoding analysis from 0 to 10.35 sec in steps of the TR (450 ms). Thus, instead of assuming a fixed temporal profile, decoding time courses were fitted with a single-gamma response function to determine amplitude and temporal delay (*see Materials and Methods*). For comparison, the same analysis was also performed for V1.

Our results reveal a dissociation between V1 representations on the one hand and anterior and posterior hippocampal representations on the other hand. As expected, V1 representations during invalid trials were dominated by the bottom-up visual input. Specifically, V1 representations were biased toward the presented (less expected) sequence compared to the expected (but not presented) sequence (**Figure 4B**). Note however, that while V1 representations generally appear to be biased toward the presented sequence, analysis of RFs covering the non-stimulated sequence RFs showed that V1 additionally represents the predicted sequence. Namely, BOLD activity at the expected (and not presented) RFs during invalid trials was found to be greater compared to BOLD activity at the not expected (and not presented) RFs during valid trials (t(27) = −2.22, p = 0.035), suggesting that V1 also represents the expected sequence in addition to the presented sequence in the probabilistic context. In contrast, in the random context the difference between valid and invalid trials was not significant t(27) = 1.18, p = 0.248), resulting in a significant Validity [Valid, Invalid] x Context [Probabilistic, Random] interaction (F(1,27) = 5.42, p = 0.028, η^2^ = 0.059).

**Figure 4.**
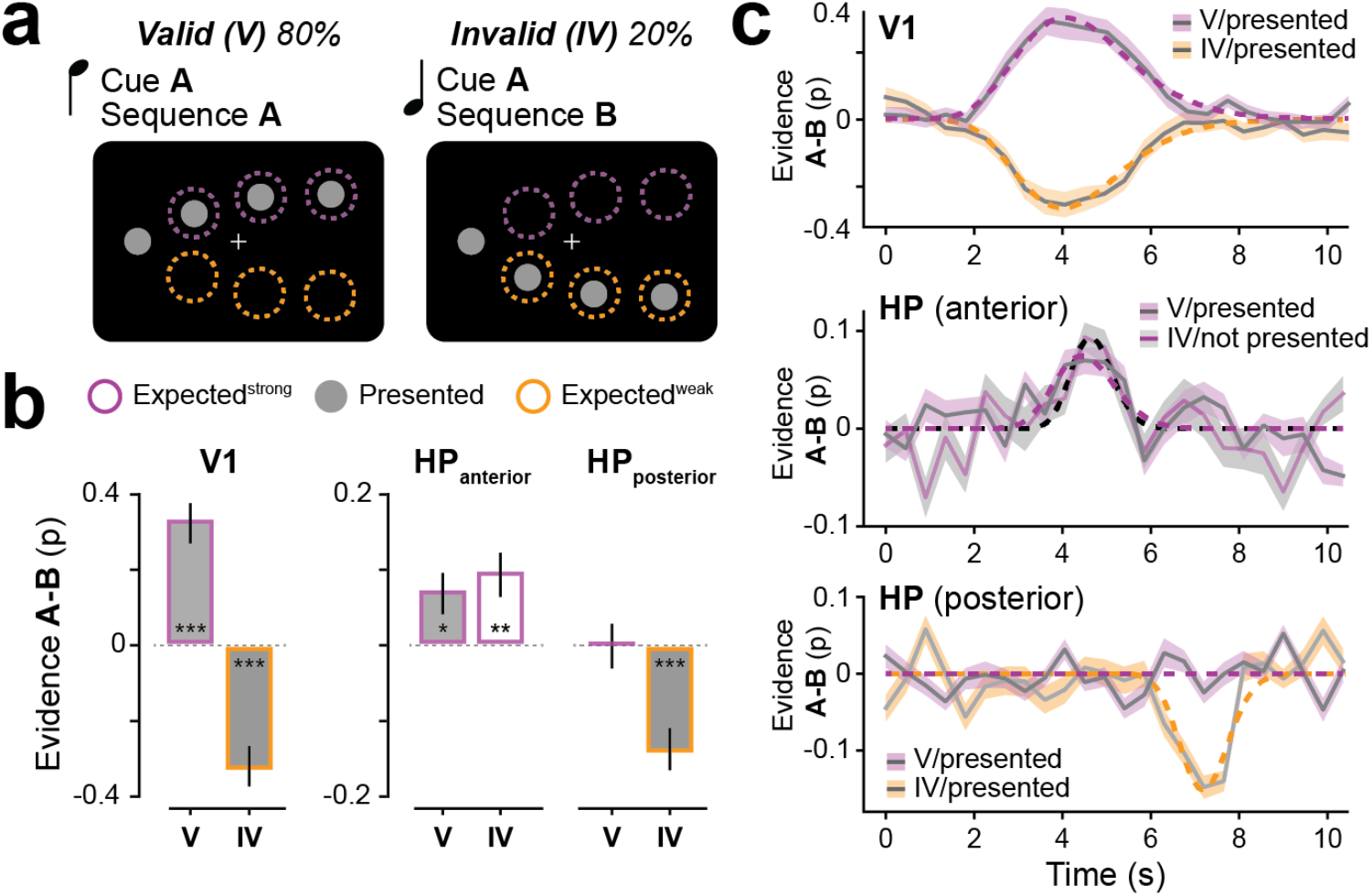
Distinct representation of stimulus and prediction. (**a**). Comparing cue valid (V) and cue invalid (IV) full sequence trials enabled us to disentangle the influence of sequence presentation and prediction. During valid trials, the presented sequence (grey) is also strongly expected (purple), while during cue invalid trials the presented sequence is less expected (orange). (**b**) Classifier evidence shows that V1 representations are biased toward the sequence presentation during both valid and invalid trials. Representations in the anterior hippocampus (HP) represent the predicted sequence during valid and invalid trials, irrespective of the presented sequence. The posterior hippocampus showed a biased representation of the presented (and less expected) sequence during invalid trials (**c**). Group averaged (N=28) decoding time courses for V1 (top), anterior hippocampus (middle) and posterior hippocampus (bottom). Dashed line denotes single-gamma fit to determine decoding amplitude and delay. Error bars denote ± s.e.m.; **P<0.001; **P<0.01; *P<0.05.

The anterior hippocampus activity patterns preferably represented the predicted sequence, irrespective of the presented sequence (**Figure 4B** middle), while representations in the posterior hippocampus were biased toward the presented sequence. Importantly, the posterior hippocampus evidence was only present during invalid trials, but not during valid trials (**Figure 4B** right). The Validity x hippocampal subpart [anterior, posterior] interaction was found to be significant (F(1,27) = 9.685, p = 0.004, η^2^ = 0.061) confirming the differential involvement of anterior and posterior hippocampus for representing predictions.

Decoding time courses revealed a V1 peak at around 4.1 s (**Figure 4C**), in line with known V1 BOLD hemodynamics (Boynton et al., 1996). Representational time-courses in the anterior hippocampus peaked at around 4.5 s, while the decoding time course of the posterior hippocampus peaked notably later at 7.1 s.

## Discussion

In this study, we sought to answer the question whether human early visual cortex (V1) and hippocampus represent multiple future expected states, with activity proportional to the probability of occurrence in the environment. Our results revealed that V1 and hippocampus showed coordinated cue-triggered forward replay that reactivates stimulus representation of expected future events. Importantly, both regions accurately represent the underlying probabilistic cue-sequence relationship of the environment by more strongly encoding the more like sequence trajectory over the less likely sequence trajectory. Further, a context manipulation in which we changed the probabilistic cue-sequence relationship, highlights the flexibility of memory-guided replay to dynamically adapt to structural changes in the environment.

Participants learned the probabilistic association of an auditory tone with a moving dot stimulus sequence that followed either an upward or downward trajectory. We studied expectations of future states by occasionally only flashing the sequence starting point together with the tone that indicated the probability of which sequence was most likely to appear. This created a situation in which the next part of the sequence was expected, but not physically shown to the participant. A multivariate pattern analysis revealed that the partial presentation was sufficient to trigger reactivation of full sequence representations in both V1 and hippocampus even in the absence of visual input (i.e., only the starting point). Importantly, correlation of anticipatory sequence representations in V1 and hippocampus revealed that forward replay between these two regions was coordinated.

### Mnemonic representations guide predictions of future events

Our results support the notion that predictions are guided by mnemonic representations of previous experiences. The hippocampus has been suggested to play a pivotal role in acquiring these regularities. One research line that is often used to probe hippocampal representations of the environmental regularities is statistical learning, where participants are presented with a constant stream of stimuli. These studies have shown that the hippocampus formation enables us to represent the structure of the environment (Kourtzi & Welchman, 2019; Schapiro et al., 2012; Sherman et al., 2020; Turk-Browne, 2019), that hippocampus can learn arbitrary relationships across different stimuli (Cohen & Eichenbaum, 1993; Davachi, 2006; Hsieh et al., 2014; Kok & Turk-Browne, 2018), and that representations of such relationships are heavily impaired after hippocampal damage (Chun & Phelps, 1999; Finnie et al., 2021; Jadhav et al., 2012; Schapiro et al., 2014; Sutherland et al., 1989).

Especially relevant in the context of the present study is that hippocampus has also been involved in exploiting learned regularities after learning is complete (Stachenfeld et al., 2017). Specifically, one of the prominent hippocampus functions is the ability to retrieve associated item information based on partial information, a function called pattern completion (Deuker et al., 2013; Leutgeb & Leutgeb, 2007; Treves & Rolls, 1994). Pattern completion has been mostly studied during episodic memory recall, but is also suited for perceptual predictions based on contextual cues (Barron et al., 2020; Bosch et al., 2014; Eichenbaum & Fortin, 2009; Hindy et al., 2016; Kok & Turk-Browne, 2018). In fact we recently argued that this hippocampal mechanism might drive the reinstatement of full stimulus sequences that we observed previously in V1 (Ekman et al., 2017). Computationally, pattern completion is thought to be implemented via recurrent, auto-associative fibers in the CA3 hippocampal subfield (Carr et al., 2011; Treves & Rolls, 1994), from where the retrieved information is output to CA1 and further downstream via bidirectional connections to sensory cortex (Eichenbaum et al., 2007; Lavenex & Amaral, 2000).

### Reinstated sensory details encode the probability of future events

In the context of a theoretical framework whereby hippocampus represents a generative model of the environment (Stachenfeld et al., 2017) and feeds that information to V1, it appears striking that V1 encodes representations of both the strongly and weakly predicted sequence concurrently, instead of reinstating only a single prediction of the most likely sequence. This result is in line with Bayesian theories of neural coding that propose that populations of neurons represent stimuli as probability distribution (Pouget et al., 2013) to deal with ambiguous and uncertain information. Previous studies have shown that human observers employ knowledge of uncertainty when making perceptual decisions and that probability distributions reflecting sensory uncertainty can be extracted from human visual cortex (van Bergen et al., 2015). Our results show that the probability of the respective sequence was proportional to the BOLD amplitude at receptive field locations corresponding to the expected stimulus. It is however noticeable that anticipated BOLD amplitude did not scale with probability across different scanning sessions. If that were the case, one would expect that anticipated activity for the less likely sequence in the probabilistic session (20% likely) should be lower compared to the anticipated activity for the equally likely sequence in the random session (50% likely). Our results show that this was however not the case, potentially indicating that sequence probability is not encoded in absolute terms, but rather scaled by the relative context.

An advantage of our paradigm using non-overlapping moving dot sequences is that it allowed us to precisely quantify reactivated stimulus representations by characterizing activity at the receptive field level. In contrast, studies looking at sensory reinstatement through the window of representational pattern analysis cannot directly quantify the perceptual detail with which perceptual representations are reinstated for centrally presented stimulus sequences. Our analysis showed that the same receptive fields that respond to the visual sequence, are also reactivated during reinstatement, confirming the low-level, perception-like properties of replay.

Given that hippocampal representations have predominantly been studied in the context of spatial navigation, it might be surprising to find that hippocampus also represents visual sequences in the context of our paradigm. However, based on recent reports a picture emerges in which hippocampal representations constitute a more general mechanism for encoding also non-spatial continuous variables (Stachenfeld et al., 2017) including tone frequency (Aronov et al., 2017) or abstract knowledge (Constantinescu et al., 2016). Our findings add to a growing body of literature that indicates that the hippocampus can provide predictive representations beyond navigation.

### Spatiotemporal dissociation of hippocampal representations

Our results reveal an interesting distinction between the anterior and posterior hippocampus. In line with earlier work (Hindy et al., 2016; Zeidman & Maguire, 2016), the anterior part of the hippocampus represented the prediction of the upcoming sequence, while the posterior part of the hippocampus showed a strong prediction error-like response in case expectations were violated. It is possible that the prediction error response reflects the ongoing mechanism by which the hippocampus learns regularities. In fact, Bein et al. 2020 have recently shown that prediction errors put the hippocampus in an encoding mode and shift the balance from mnemonic retrieval to sensory processing, effectively updating the model of the world.

In absence of a fine-grained retinotopic organization like in V1, probing hippocampal representations is challenging. Similar to other studies (Hindy et al., 2016; Kok & Turk-Browne, 2018), we used a multivariate pattern analysis approach that does not assume a specific hippocampus encoding schema. However, given that the underlying mechanisms for hippocampal representations of visual space remain unknown, it is possible that instead of forming a perceptual representation, hippocampus might rather encode a more abstract, temporally structured representation by simply indexing discrete stimulus features stored in V1 (Teyler & DiScenna, 1986). It remains therefore an interesting avenue for future studies to reveal how hippocampus encodes visual representations. For instance, Knapen (2021) and Silson et al. (2021) have recently used receptive field mapping to show that the anterior hippocampus features a basic visual field representation. It is therefore possible that such subtle receptive field biases might contribute to the encoding of visual sequences that were used in the present study.

One limitation of our study is that due to the low temporal resolution of BOLD fMRI we cannot exclude the possibility that the simultaneous prediction of both sequences actually appears sequentially, potentially starting with the more likely sequence, and not in parallel. Such sequential hippocampal replay has been observed in the context of spatial navigation in rodents (Gupta et al., 2010; Johnson & Redish, 2007). We think that the novelty of the current study is to show the presence of probabilistic replay in humans, while questions pertaining to the temporal order of events might be better suited for future studies using other modalities like magnetoencephalography (Kurth-Nelson et al., 2016; Liu et al., 2019).

Perception depends on both the current sensory input and on previous experience. The current study provides novel insights into how the probability of future events is encoded in predictive representations. Our results highlight the tight link between sensory manifestation of predictions in early visual cortex and associative representations in the hippocampus.

## Material and Methods

### Data and code availability

All data and code used for stimulus presentation and analysis is available on the Donders Repository https://data.donders.ru.nl/[direct link to dataset]. [*Note that during the review process, reviewers have anonymous access using the following link [retracted]. After publication this link will be made publicly available*.]

### Participants

Twenty-eight right-handed participants (16 female, mean age = 25 years) were recruited from the student population at the Radboud University in Nijmegen and invited for two separate fMRI sessions. Sample size was decided on prior to the collection of data and was aimed at being able to detect experimental effects that had at least moderate effect size (Cohen’s d>0.6). Participants gave written informed consent in accordance with the institutional guidelines of the local ethical committee (CMO region Arnhem-Nijmegen, The Netherlands) and received monetary compensation for their participation. One participant completed only one session. All participants had normal or corrected-to-normal visual acuity.

### MRI acquisition

Functional and anatomical images were acquired using a 3T Prisma MRI system (Siemens, Erlangen, Germany) equipped with a 32-channel head coil. Each of two MRI sessions lasted approximately 2 h, during which we acquired (*i*) a T1-weighted anatomical scan, (*ii*) functional scans to measure BOLD activity during the experimental paradigm (*iii*) wholebrain functional scans to improve co-registration of the functional sequence, and (*iv*) functional scans to perform retinotopic mapping.

The retinotopic mapping was only performed once, either during the first or during the second session. BOLD activity for the retinotopic mapping was measured using a T2*-weighted gradient-echo EPI sequence (TR/TE = 1800/30 ms, 26 transversal slices, voxel size 2×2×2 mm, 60° flip angle). BOLD activity for the experimental runs was measured using a T2*-weighted multiband sequence (acceleration factor (MB) = 3; TR/TE = 450/39 ms, 15 transversal slices, voxel size 2.4×2.4×2.4 mm, 45° flip angle; slice gap = 10 %). Slices were carefully positioned to cover the relevant parts of primary visual cortex and hippocampus (**Figure 3B**). Anatomical images were acquired with a T1-weighted MP-RAGE sequence (TR/TE = 2300/3.03 ms, voxel size 0.8×0.8×0.8 mm, 8° flip angle).

### Stimuli

Visual stimuli were rear projected on a screen (luminance-calibrated EIKI projector, 1,024 × 768 resolution, 60 Hz refresh rate) located 86.6 cm from the participant’s eyes at the head of the scanner table. The screen was viewed using a mirror attached to the headcoil. We presented one of two moving dot sequence consisting of four white dots on a black background (spatial coordinates sequence up: x=[−6°, −2°, 2°, 6°]; y=[0°, 2.1°, 3.43°, 4°]; spatial coordinates sequence down: x=[−6°, −2°, 2°, 6°]; y=[0°, −2.1°, −3.43°, −4°]). Noticeably, the first dot location was identical for the two spatial sequences. Each dot had a diameter of 0.8° and was shown for 34 ms followed by a blank screen of 17 ms. Occasionally, on 15.6% of all trials the last dot of the sequence was shown after a blank screen of 170 ms. The total duration of the moving dot sequence amounts to 187 ms (with 17 ms ISI), or 340 ms (with 170 ms ISI). A fixation-cross (0.7°) was shown at the center of the screen and participants were instructed to maintain fixation throughout the experiment. The auditory cue (pure tone, 700 or 1,400 Hz) was presented over MR-compatible earphones for a duration of 100 ms. Between the offset of the auditory cue and the onset of the visual sequence was a delay of 200 ms.

### Experimental Design

Participants were invited for two separate fMRI scanning sessions (probabilistic and random context) that were on average 14 days apart. The order of the sessions was counterbalanced across subjects. Both sessions contained the same task but differed in the probabilistic relationship between the auditory cue and the visual sequence. In the probabilistic session, the auditory cue predicted the visual sequence in ~80% (79.2%) of the trials. Specifically, the high-tone (cue A) was more likely to be followed by the upward sequence (sequence A) and the low-tone (cue B) was more likely to be followed by the downward sequence (sequence B). In the non-predictive session, the relationship between the auditory cue and the visual sequence was at chance level (50%). Participants were explicitly instructed about the cue validity in the respective session.

Each session consisted of two parts, an initial learning period and a task period. During the learning period only full sequence trials were shown, containing the auditory cue (A, B) and 4 dots of the respective sequence (A, B). Participants were instructed to detect irregular trials (occurrence 15.6%), in which the last dot in the sequence was presented after 170 ms, instead of 17 ms as in regular trials. Participants had to report on each trial whether a sequence was regular (left button press, right index finger), or irregular (right button press, right middle finger). Feedback was given on each trial for 200 ms in form of a color change of the fixation cross (green: correct; red: incorrect/miss). Additionally, a summary screen was presented at the end of each learning block informing participants about the percentage of correct responses. Learning trials were separated by a fixed ITI of 450 ms.

The learning part consisted of three blocks. In order to facilitate learning, the first and second learning block (96 trials, ~4 min each) featured only one of the two auditory cues, respectively. For instance, one participant would start with a learning block featuring only cue A, followed by sequence A, or sequence B depending on the context (probabilistic vs random). After that, in the second learning block, the participant would be presented only with cue B, followed by sequence A, or sequence B. The third and final learning block contained both cue A and cue B trials (192 trials, ~8 min.). The order of learning block one and two was randomized such that half the participants started with the cue A learning block. Learning blocks were separated by 3 minutes of rest. Learning block one and two contained two null-events (duration 10.8 s) during which only the fixation-cross was shown. Learning block three contained 5 null-events. In sum, during learning participants were exposed to 384 visual sequence trials in total, with 304/80 valid/invalid trials in the probabilistic session and 192/192 valid/invalid trials in the random session.

During the task period, participants were also presented with auditory cues and visual sequences, but were instructed to perform a detection task at fixation. On every trial (duration 13.95 s, 31 volumes), the fixation cross was dimmed once for a brief moment (minus 60% contrast for 17 ms) and participants had to press a button with the right index finger. The timing of the dimming occurred in the temporal range between 450 ms and 13950 ms, in steps of 900 ms, randomly sampled from a uniform distribution (without replacement). A summary of participants performance was given at the end of each block.

The task period consisted of 2 blocks, each with 52 trials and 4 null-events (10.8 s). 16 out of 52 trials per block were replay trials in which only the auditory cue (high or low) and the starting point of the visual sequence was shown. In total, the task period contained 104 trials with 32 replay trials (~30%) and 72 sequence trials (~70%). No odd trials were shown during the task period. Trial order was pseudo-randomized to ensure no repetitions of replay trials. Further, task blocks never started with a replay trial.

### pRF estimation

The data from the moving bar runs were used to estimate the population receptive field (pRF) of each voxel in the functional volumes using MrVista (http://white.stanford.edu/software). In this analysis, a predicted BOLD signal is calculated from the known stimulus parameters and a model of the underlying neuronal population. The model of the neuronal population consisted of a two-dimensional Gaussian pRF, with parameters x0, y0, and σ0, where x0 and y0 are the coordinates of the center of the receptive field, and σ0 indicates its spread (standard deviation), or size. All parameters were stimulus-referred, and their units were degrees of visual angle. These parameters were adjusted to obtain the best possible fit of the predicted to the actual BOLD signal. This method has been shown to produce pRF size estimates that agree well with electrophysiological receptive field measurements in monkey and human visual cortex (Klink et al., 2021). For details of this procedure, see (Dumoulin & Wandell, 2008; Kay et al., 2015). Once estimated, x0 and y0 were converted to eccentricity and polar-angle measures and co-registered with the functional images using linear transformation. Only voxels with a model fit of R^2^ ≥ 5% were considered.

### ROI selection

V1 and hippocampus region of interests (ROIs) were determined using the automatic cortical parcellation provided by Freesurfer (Fischl, 2012) based on individual T1 images. Anatomical V1 and hippocampus masks were then transformed into native space using linear transformation. The anatomical V1 mask was restricted to voxels with receptive fields size ≤ 2.5 dva to increase the spatial specificity of a voxels’ response. ROIs for the retinotopic locations corresponding to the sequence dot locations were chosen in the following way. First, for each of the 7 dot locations (e.g. starting dot sequence A/B), all voxel with a receptive field covering that respective location were selected. However, this could lead to an unequal number of selected voxels between dot locations. Second, in order to balance the number for voxel across dot locations, for each participant, we determined the number of voxels with a receptive field of the starting dot location (i.e., x=-6°, y=0°). In case the other dot locations were covered by fewer voxel receptive fields, we reduced the number of voxels for each dot ROI by decreasing the number of voxels until the number was equal to the starting dot location. Voxels were removed based on their statistical z-value in the independent learning block (stimulation vs baseline contrast). Voxels with lower values were removed first. An additional, nonstimulated control ROI was chosen that mirrors the starting dot location at x=6°, y=0° using the same procedure.

In order to exclude the possibility of signal spillover from the nearby lateral geniculate nucleus (LGN), a subcortical thalamus mask (including the LGN; (Collins et al., 1995)) was removed from the hippocampus ROI. Anterior and posterior hippocampus ROIs were created by following the procedure outlined in Silson et al. (2021). In short, hippocampus voxel indices were sorted by the y-axis, which codes for cortical anterior-posterior position. These indices were then separated into two approximately equally sized ROIs.

### fMRI preprocessing

Images were preprocessed using FSL (Smith et al., 2004) including motion correction using six-parameter affine transform, temporal high-pass filtering (128 s), and spatial smoothing with a Gaussian kernel (full width at half maximum of 5 mm). All analyses were carried out in native subject-space. The first 10 volumes of each run were discarded to allow for signal stabilization.

### BOLD amplitude modulation

A general linear model (GLM) was used to fit individual BOLD responses and obtain estimates of signal change per voxel. The GLM consisted of 6 regressors of interest: full sequence (cue A/B, valid, invalid), replay (cue A/B), one regressor of no interest modeling the task instructions and performance summary screen and 24 motion regressors (‘standard+extended’ option in FSL FEAT, representing the 6 estimated realignment parameters, its 6 derivatives plus the 12 corresponding squared regressors of the former regressors). Parameter estimates were estimated for each run separately and averaged within each ROI. In order to control for stimulus unspecific BOLD modulations, we subtracted the PE of a non-stimulated control ROI (see ROI selection) from the PEs of the stimulus ROIs. Finally, PE’s were compared across participants using repeated-measures ANOVA.

### Decoding analysis

The decoding analysis was performed with scikit-learn (Pedregosa et al., 2011). Individual voxel time courses were low-pass filtered using a Savitzky-Golay filter with a window length of 5 TRs and polynomial order of 2 (Savitzky & Golay, 1964) and normalized to z-scores. A logistic regression classifier (default values, L2 regularization; C=1) was trained to distinguish between sequences A and B during valid full sequence trials. In addition to the two classes of interest a third class of no interest (null event, fixation cross only) was included during classifier training. Similar to a covariate in a GLM, this class might capture noise contributions that present throughout the experiment (e.g. from respiration) and should not be used to distinguish between the classes of interest. The classifier training was performed on single trials from one run. Classifier training and testing was always performed across runs to ensure independent datasets (i.e. training on run 1 and testing on run 2, then training on run 2 and testing on run 1). The trained classifier was then applied to replay trials (or invalid full sequence trials). Before applying the trained classifier, all trials of a respective condition were averaged to maximize signal-to-noise ratio. The classifier output consists of a probability for each class (i.e., sequence A, B, rest; probabilities sum up to 1), which were averaged across independent runs and tested across participants and conditions using a rm-ANOVA.

For the time-resolved decoding analysis, the described steps were repeated for each TR from 0 to 10.35 s. For the standard decoding analysis, we chose an average of 3 volumes corresponding to the expected hemodynamic peak at around 4.05-4.95 s. A Searchlight analysis was performed repeating the same decoding analysis within a sphere of r=6 mm.

### V1-hippocampus correlation

Across subject correlation analysis was performed to probe coordinated replay in V1 and hippocampus. For each participant, classifier evidence values represent the probability that the cued sequence was replayed minus the probability of the noncued sequence (i.e. probability sequence replay A minus B during cue A trials and probability sequence replay B minus A during cue B trials). Classifier evidence for V1 and hippocampus were then tested for dependency using Spearman’s rank correlation coefficient, separately for the probabilistic and random session. Differences in Spearman correlation coefficients between the probabilistic and random session were tested using a non-parametric test with 10.000 permutations.

### Control for eye movements

Participants were instructed to maintain fixation throughout the whole experiment. Eye positions were recorded with a video camera at 50 Hz sampling rate under infrared illumination (Eye Track-LR camera unit, SMI, SensoMotoric Instruments). Eyeblink artifacts were identified by differentiating the signal to detect eye pupil changes occurring too rapidly (< 60 ms) to represent actual dilation. Blinks and samples in which the corneal reflection was not reliably detected were removed from the signal using linear interpolation. Eyetracking data was gathered in the scanner for 21 of the 28 participants during the probabilistic session and for 24 of the 27 participants for the random session. We calculated the mean gaze as a function of the four stimulus locations and task conditions. Mean horizontal gaze position did not vary with stimulus position, neither for the probabilistic session (ANOVA, *P* = 0.56) nor for the random session (ANOVA, *P* = 0.72).

## Acknowledgments

This study was supported by the Netherlands Organization for Scientific Research (Veni Grant No. 016.Veni.195.435), awarded to M.E. and the James S. McDonnell Foundation (JSMF Scholar Award for Understanding Human Cognition), the European Union Horizon 2020 Program (ERC Starting Grant 678286, “Contextvision”), awarded to F.P.d.L. We thank Eva Berlot and Britta U. Westner for comments on an earlier version of the manuscript.

## Author contributions

M.E., and F.P.d.L., conceived and designed the experiments. M.E. and G.G. collected the data, M.E. conducted the data analyses. M.E., G.G. and F.P.d.L. wrote the manuscript.

